# Temperature variation drives coordinated scaling of temporal and dynamic features of transcription in embryonic development

**DOI:** 10.1101/2023.08.09.552666

**Authors:** Gabriella Martini, Hernan Garcia

## Abstract

Temperature affects the timing of development in most poikilothermic organisms that cannot regulate their internal body temperature. In the fruit fly, *Drosophila melanogaster*, similar quantitative trends characterize changes in the timing of morphological events in embryogenesis from cellularization to hatching across a 10°C temperature range, such that the relative duration of each of these stages is temperature-independent. However, the extent to which the timing of the individual molecular and cellular processes underlying these morphological events recapitulates this relationship with temperature is largely unknown. Here, we characterized how the spatiotemporal dynamics of the process of transcription, which are so fundamental to cell fate commitment, scale with temperature in single cells of living fly embryos. Using the *hunchback* gene as a case study, we discovered that the duration of the cell cycle and the temporal and dynamic features of *hunchback* transcription scaled in a coordinated fashion such that the relative rates of all observed processes were temperature independent and, perhaps most surprisingly, such that the total amount of mRNA produced by the gene is unaltered by temperature changes. Our approach provides a crucial tool for understanding both developmental robustness in the face of environmental variation and for applying biochemical approaches in living *Drosophila* embryos.

## Introduction

Most organisms must be able to adapt to new climates and contend with large ranges in environmental temperature. Temperature variation affects phenomena ranging from development, to physiology, to behavior, especially in ectothermic organisms that do not consistently regulate their body temperature. Embryonic development, which involves a plethora of highly dynamic biochemical processes underlying cellular decision making and morphological changes, poses a particularly intriguing challenge for understanding how body plans remain robust in the face of significant temperature variation. For example, the developmental program of the fruit fly, *Drosophila melanogaster*, remains largely indifferent to temperature variation (Kuntz and Eisen (2014); Chong et al. (2018)), even when subject to temperature gradients across the embryo (Lucchetta et al. (2005)). Similarly, even though the dynamics of vertebrate segmentation have been shown to be highly dependent on temperature (Schroter et al. (2008)), the segmentation of fish, for example medaka, is indifferent to the modulation of temperature between 15°C and 35°C (Vibe (2021)).

While developmental outcomes appear unchanged over a wide range of temperatures, the rates of the biochemical processes underlying embryonic development are expected to vary by drastically different amounts (Section). Because developmental dynamics rely on the co-ordination of various biochemical processes, it is not clear how the same developmental outcome can be achieved if the relative rates at which these underlying biochemical processes proceed are temperature-dependent (Falahati et al. (2021)).

Recent work in the fruit fly, *Drosophila melanogaster*, has shed light on the relationship between the timing of developmental processes and temperature at various physical scales. At the macroscopic level, it has been observed that the timing of the appearance of the vast majority of morphological features throughout development, some of which are shown schematically in Figure 1A, scaled with a single time constant relating each pair of temperatures (Kuntz and Eisen (2014); Crapse et al. (2021)). Specifically, consider the temperature dependence of the timing of morphological events outlined schematically in Figure 1B. Suppose that, at 25°C, which we will use as a reference temperature throughout this work, a given developmental event, *n*, occurs at time *t*_*n*_(*T* = 25°C) after fertilization. Generalizing to any temperature, *T*, at which *Drosophila* embryogenesis proceeds healthily, we can write the time taken to reach this same event following fertilization, as *t*_*n*_(*T*). Kuntz and Eisen (2014) showed that the scaling of this time with temperature is given by the same temperature-dependent factor *α*(*T*), shown schematically in Figure 1C, for all developmental features examined. As a result, the time to reach any point in development at temperature *T* is given by

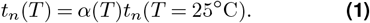

**Figure 1.**
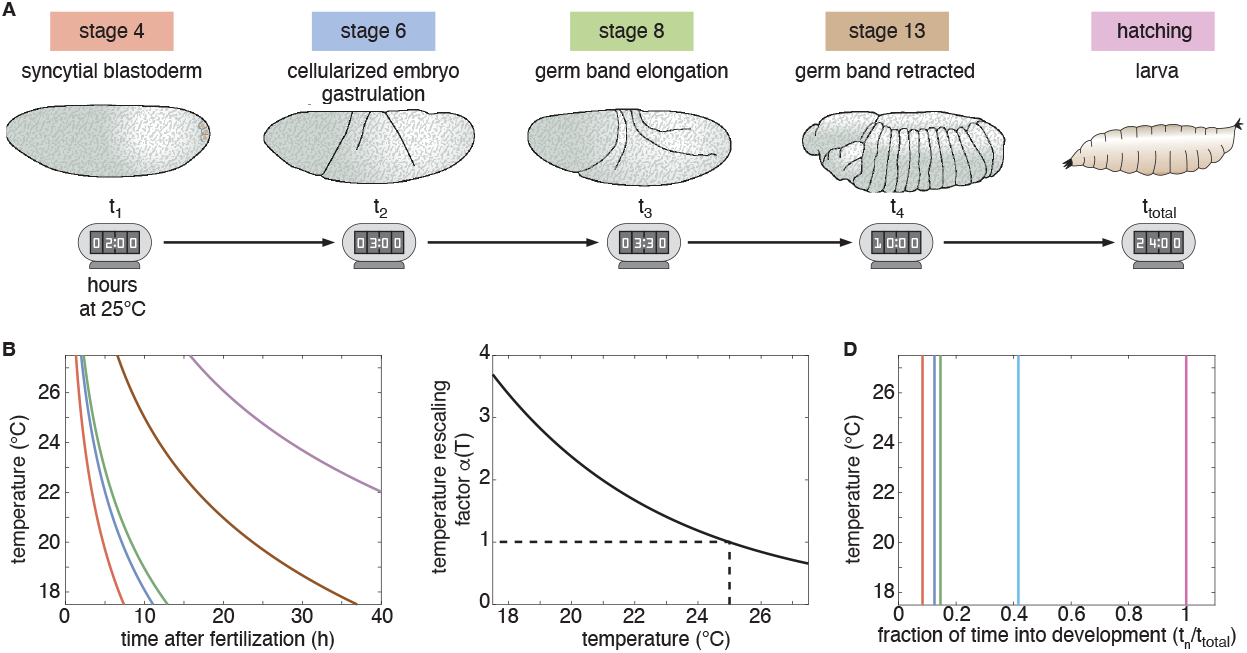
Schematic of unified temperature scaling in development. (A) Timing of the appearance of a number of developmental landmarks in *Drosophila melanogaster* embryogenesis at 25°C. (B) Illustrative dependence of the timing of developmental landmarks with temperature. The colored lines corresponds to the morphological events schematized in (A). The dashed line indicates the timing of events at 25°C as in (A). (C) Universal scaling factor *α*(*T*) dictating the pace of development as a function of temperature. Note that, as defined here, *α*(*T* = 25°C) = 1 such that 25°C becomes a reference temperature. (D) When time is rescaled by the total time to reach the final developmental landmark, all morphological events occur at the same fractional time into development independently of temperature.

In other words, consider the total time of development, spanning from fertilization to larval hatching, given by *t*_*total*_(*T*) = *α*(*T*)*t*_*total*_(*T* = 25°C). If we normalize the time, *t*_*n*_(*T*) at which developmental event, *n*, occurs by *t*_*total*_(*T*), we see that each morphological event takes place at a consistent proportion of the total time of development, as shown schematically in Figure 1D. This temperature independence in the relative timing of morphological events could suggest the existence of a kind of “developmental clock” that uniformly scales the pace of development with temperature (Kuntz and Eisen (2014)). But how might such a clock work?

The physical scale at which the developmental clock operates remains an open question. Specifically, does this developmental clock only determine the pace of morphological events, or does it also dictate the timing of the underlying molecular processes? A recent work in the fruit fly examined the temperature dependence of the cell cycle that drives rhythmic mitotic divisions in early development (Falahati et al. (2021)). The authors found that, with the exception of prophase, all remaining stages of the cycle—interphase, prometaphase, metaphase, anaphase, telophase—displayed a comparable scaling with temperature, once again consistent with Equation 1 (Falahati et al. (2021)).

While the timing of mitosis seems to be mostly consistent with the existence of a developmental clock, the temperature dependence of other molecular processes key to embryonic development, such as the transcriptionally-mediated establishment of the differentiated gene expression patterns specifying the body plan of the adult fly, has not been assayed in the fruit fly to date. Thus, we wanted to understand whether and how transcription compensates for temperature-driven changes in the timing of development. Specifically, do the dynamics of the plethora of processes underlying transcriptional—i.e. the time window during which transcription occurs within the cell cycle and the rates of transcriptional initiation, elongation and termination—all display comparable temperature scaling *α*(*T*) with respect to each other, as well as with respect to the morphologically defined developmental clock?

In this work, we set out to determine how the timing of transcriptional activity and the rates that define the main steps of the transcription cycle scale with temperature using the early fly embryo as a case study. We used the MS2 system to image the transcriptional dynamics of one of the most studied gene expression patterns in all of animal development: the step-like expression pattern of the *hunchback* gene (Margolis et al. (1995); Driever et al. (1989); Perry et al. (2012); Park et al. (2019)). As expected, we find that the rates of transcriptional initiation, elongation, and termination increase with temperature. Yet, at the same time, the length of the cell cycle contracts by exactly the right amount such that the total amount of mRNA produced—the product of the initiation rate and the window of time over which transcription occurs during a nuclear cycle—remains unchanged over a temperature range of 10°C. Thus, we discovered a compensated scaling of transcriptional activity and the window over which it occurs, which ensures a conserved molecular quantity over a large temperature range.

Finally, in an attempt to resolve the temperature dependence of the mean transcription initiation rate at an even finer scale, we examined how temperature affects the transcriptional bursting dynamics that seem to underlie the transcription of most genes in early development (Lammers et al. (2020b)). We find that the temperature scaling of the mean rate of transcriptional initiation is driven by coordinated scaling of the burst amplitude, duration and frequency. Thus, we discovered that the spatiotemporal dynamics of every transcriptionally-associated process within our experimental reach shares the same quantitative temperature scaling *α*(*T*). As a result, we speculate that coordinated temperature scaling in development goes beyond the timing of macroscopic morphological events and of the microscopic processes driving most phases of the cell cycle, to the dynamics of transcription, which dictates the adoption of cellular fate throughout development.

## Results

### An experimental platform to study the temperature dependence of transcriptional dynamics in living embryos

To study the temperature dependence of transcriptional control in the early fly embryo, we dissected the transcriptional dynamics of the *hunchback* gene, one of the key players in the gene regulatory network that dictates body plan segmentation (Margolis et al. (1995); Perry et al. (2012)). During early development, as zygotic transcription is just beginning and the embryo is in the syncytial blastoderm stage, *hunchback* expression is primarily regulated by the maternally deposited Bicoid activator (Driever and Nusslein-Volhard (1989); Park et al. (2019)). The resulting expression pattern is an anterior accumulation of *hunchback* mRNA with a sharp, steplike decline along the middle of the embryo’s anterior-posterior axis that is clearly observable during the 13th nuclear cycle, starting at about an hour and 40 minutes into development at 25°C. This step sharpens considerably around 2 hours into development at 25°C during the 14th nuclear cycle, in which the first cellular membrane cleavages occur. (Figure 2A; Gregor et al. (2007); Bothma et al. (2015); Perry et al. (2012)).

**Figure 2.**
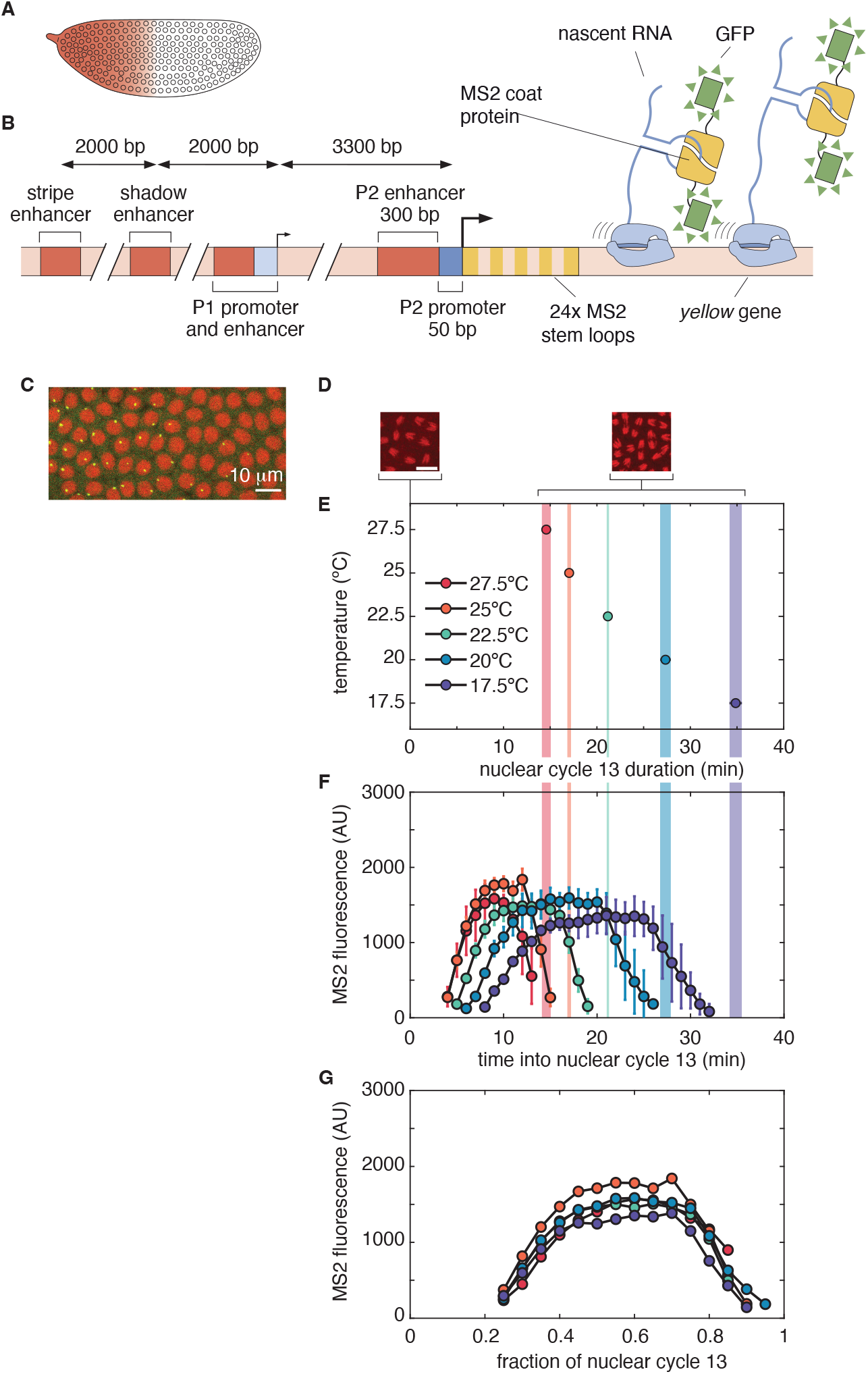
Temperature control and simultaneous monitoring of transcriptional activity in the early fruit fly embryo. (A) Cartoon of the step-like *hunchback* gene expression pattern that constitutes the basis of this work. (B) A *hunchback* BAC reporter is labeled with the MS2 system. Here, 24 repeats of the MS2 sequence are incorporated in the 5’ end of the reporter gene. When transcribed, these sequences form stem loops that are recognized by maternally deposited MS2 coat protein (MCP) fused to GFP. (C) Sites of nascent transcription emerge as fluorescent puncta (green), while nuclear divisions can be tracked using a fusion of Histone to RFP (red). (D) Measurement of the length of nuclear cycle 13 by quantifying the time between subsequent anaphases. Scale bar is 10 *µ*m. (E) Duration of nuclear cycle 13 as a function of temperature. The number of embryos included in the calculation are 5, 12, 6, 7, and 4 going from 27.5°C to 17.5°C, respectively. Embryo-specific calculations of the mean duration of the 13th nuclear cycle included 62–112 individual nuclei. The duration of nuclear cycle 13 was then calculated by taking the mean over all embryos developing at a given temperature. Errorbars are standard errors calculated over embryos. (F) Mean MS2 fluorescence intensities at 30% embryo length as functions of time into nuclear cycle 13. Fluorescence traces were averaged over all nuclei with detectable levels of transcription located at 30% *±* 1.25% of the embryo length. Fluorescence signals were rescaled to account for temperature-dependence in the GFP signal (see Materials and Methods). Errorbars are standard errors of the mean fluorescence intensity traces calculated over embryos. The number of embryos included in the calculation are 2, 7, 5, 3, and 2, going from 27.5°C to 17.5°C. (G) Mean MS2 fluorescence intensities as a functions of the fraction of nuclear cycle 13. Errorbars are omitted for visual clarity, but are the same as those shown in panel (F).

To observe *hunchback* transcriptional activity *in vivo*, we used a previously characterized *hunchback* BAC reporter labeled with an MS2 stem loop cassette at its 5^*′*^ end (Bertrand et al. (1998); Garcia et al. (2013); Bothma et al. (2015)). This reporter construct recapitulates the most salient features of the endogenous gene expression pattern (Bothma et al. (2015)). When the MS2 sequence located in the reporter is transcribed, the sequence folds to create stem loops that bind with the maternally provided MS2 coat protein (MCP) fused to eGFP (Figure 2B). As a result, sites of nascent transcript formation appear as fluorescent puncta (Figure 2C), with intensities that are proportional to the number of actively transcribing RNA polymerase molecules at each locus.

To systematically modulate the temperature at which embryonic development was observed, we controlled the sample temperature using a water-based microscope stage-top incubator. The temperature in the incubator was regulated using a heating and cooling water bath from which water was circulated through the base and lid of the sample chamber as well as through an objective collar, creating a “water jacket” around the samples. Sample temperatures were monitored using a fine gage thermocouple. The temperature of the room containing the laser-scanning confocal microscope was also shifted to minimize potential temperature gradients across the sample chamber and temperature fluctuations. This system yielded stable control of the chamber temperature over the 10°C range from 17.5°C to 27.5°C, with sample temperatures confirmed to be within 0.3°C of the intended temperature for each experiment using a fine gage thermocouple.

We further confirmed our ability to accurately control the sample temperature by measuring the duration of nuclear cycle 13 for embryos developing inside the incubation chamber. Specifically, we measured the time between subsequent anaphases as reported by an RFP fusion to the Histone *his2av* gene (Figure 2D). Figure 2E shows measurements of the nuclear cycle durations at 17.5°C, 20°C, 22.5°C, 25°C, and 27.5°C, the same temperatures recently assayed by Kuntz and Eisen (2014) when dissecting the timing of morphological features in fly development. Inspection of our findings reveals a monotonic decrease in nuclear cycle duration with increasing temperature, with cycle durations that are consistent with expectations based on recent experiments (Figure S2; Falahati et al. (2021)).

While the nuclear cycle is composed of a series of molecular processes, we can treat it as a single coarse-grained process with an effective rate. Under this assumption, we use the so-called *Q*_10_ to characterize the temperature dependence of this effective rate. In general, the *Q*_10_ of a reaction is an empirical measure of the ratio between reaction rates across a 10°C interval. Specifically, in this case we calculate the *Q*_10_ of the effective rate of transitioning through nuclear cycle 13 by measuring the fold-change in the length of the nuclear cycle between 17.5°C and 27.5°C to find *Q*_10_ *≈* 2.4, meaning that nuclear cycle 13 takes approximately 2.4 times as long at 17.5°C as it does at 27.5°C. This value for the cycle 13 *Q*_10_ is very similar to those observed previously for the coarse-grained effective rate of embryogenesis evaluated across the same temperature range (Kuntz and Eisen (2014)). Thus we have confirmed our capacity to control the sample temperature with an experimental setup that makes it possible to simultaneously measure transcriptional dynamics at the single cell level using the MS2 system.

### Transcriptional activity scales with the nuclear cycle length

Focusing on spatial windows of 2.5% of the embryo length, and averaging the corresponding mean MS2 fluorescence intensity traces of the *hunchback* reporter across multiple nuclei and embryos within that window reveals the dynamics of RNA polymerase molecules actively transcribing the gene. Figure 2F shows that, while the overall fluorescence levels of these signals (after correcting for temperature changes in GFP fluorescence as described in Materials and Methods) are comparable at all temperatures, the timing of the fluorescence signals varies significantly. Specifically, the time at which the signal is first detected, the time at which maximum fluorescence is reached, and the time at which the signal intensity begins decreasing all depend on temperature. Interestingly, time traces obtained at different temperatures reach near-zero fluorescence close to the end of their corresponding nuclear cycles. This cessation of transcription at the end of the nuclear cycle is consistent with previous observations suggesting that transcription is shut off as a result of entry into mitosis (Shermoen and O’Farrell (1991); Rothe et al. (1992); Gottesfeld and Forbes (1997); Parsons and Spencer (1997)).

Inspired by previous work (Kuntz and Eisen (2014)), we normalized the time into the nuclear cycle by the temperature-specific nuclear cycle durations (Figure 2E). We then considered the MS2 fluorescence (Figure 2F) as a function of the fraction of the nuclear cycle (Figure 2G) which revealed a near-complete collapse of the traces onto a single, temperature-independent profile across all assayed temperatures. Our results therefore extend previous work proposing the existence of a uniform scaling of developmental timing with temperature and show that this scaling can be used to describe the temperature dependence of the dynamics of actively transcribing RNA polymerase molecules reported by the MS2 system.

### Timing of transcriptional events scales with developmental clock

The fluorescence intensity of an MS2 spot is proportional to the number of polymerase molecules found at the corresponding locus, which is in turn determined by the combined effect of multiple steps in the transcription cycle, including initiation, elongation and termination. To zero in on which aspects of transcription are driving the temperature dependence of the MS2 signal as a function of absolute time (Figure 2F), we dissected the mean fluorescence signals to extract the rates governing the underlying biochemical processes.

To relate the fluorescence intensities as a function of time to the underlying steps of the transcription cycle, we first note that the MS2 signals shown in Figure 2F resemble a trapezoid. This trapezoidal shape can be understood in the context of a simple model of a promoter that turns on after mitosis and initiates transcription at a constant rate until shutting off again before the next mitosis (Garcia et al. (2013); Liu et al. (2021)). Specifically, consider the example shown in Figure 3A. Here, a promoter turns on at a time point *t*_*start*_, after which RNA polymerase molecules are loaded onto the gene at a constant rate *R*, yielding a linear increase in the corresponding MS2 fluorescence. After an additional time *t*_*dwell*_, the first loaded RNA polymerase molecule makes it to the end of the gene and terminates transcription. At this time point, *t*_*start*_ + *t*_*dwell*_, if the promoter is still initiating transcription, the rates at which RNA polymerases are loaded and unloaded from the reporter are equal such that the number of polymerases on the gene stays constant. As a result, the fluorescence signal plateaus until the promoter turns off at time point *t*_*end*_, after which RNA polymerases no longer initiate transcription for the remainder of the nuclear cycle. As the remaining polymerases on the reporter finish transcribing the gene and terminate transcription, the fluorescence signal decreases. Thus, the MS2 fluorescence signal contains information about the time at which transcriptional initiation starts and ends, the rate of RNA polymerase loading, and the mean dwell time of RNA polymerases on the DNA (Garcia et al. (2013); Liu et al. (2021)).

**Figure 3.**
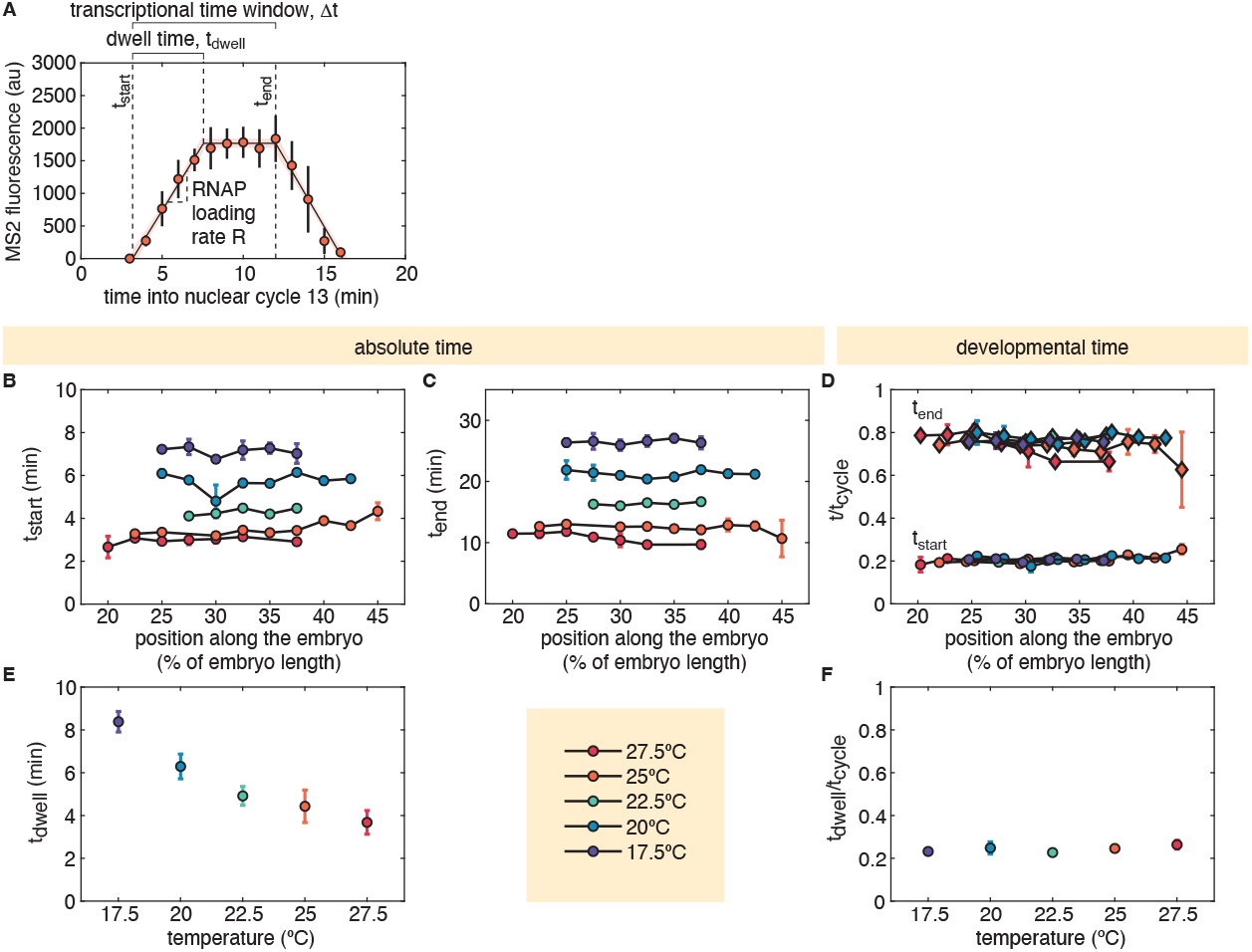
Temperature scaling of the temporal features of the transcriptional cycle captured by MS2. (A) In the context of a simple model where the promoter transcribes between times *t*_*start*_ and *t*_*end*_ by loading RNA polymerase molecules at a rate *R* that take time *t*_*dwell*_ to transcribe the gene and terminate transcription, MS2 fluorescence traces in nuclear cycle 13 are expected to resemble a trapezoid. (B-F) Inferred values for positionally-dependent parameters of the trapezoidal model of transcriptional activity. (B) Start and (C) end time of transcription as a function of the position along the embryo for different temperatures extracted from trapezoidal fits such as shown in (A). (D) When time is measured in terms of its fractional progression into the nuclear cycle, transcription starts at 20% and ends at 80% of the nuclear cycle independently of temperature. (E) RNA polymerase dwell time as a function of temperature. We assumed dwell times to be positionally independent (see Figure S3 for details) and averaged between 20-40% embryo length over multiple embryos. (F) When normalized by the duration of nuclear cycle 13 for each temperature, the dwell time becomes temperature independent. (A, average MS2 fluorescence temporal dynamics shown here corresponds to nuclei located at 30% of the embryo length averaged over 7 embryos developing at 25°C, errorbars are standard errors calculated over embryos; B-D, plotted values are the mean values calculated over parameters inferred from fits of the trapezoidal model to single embryo data, calculations included 2-5 embryos for each combination of temperature and anterior-posterior position, errorbars are standard errors over single embryo estimates; E, plotted values are the mean values calculated over positionally-independent single embryo data, temperature-dependent elongation times were calculated using data from 4-10 embryos, errorbars are standard errors over embryos.)

By fitting a trapezoid to the mean MS2 traces following the scheme presented in Figure 3A, we extracted the times corresponding to the start and end of transcription as a function of temperature and position along the embryo, as shown in Figures 3B and C. As previously reported for a minimal *hunchback* reporter construct (Garcia et al. (2013); Eck et al. (2020)), the timing of the start and end of transcription is largely independent of the position along the embryo. As expected from the temperature dependence of the nuclear cycle duration, these times decrease with increasing temperature.

Following our earlier approach introduced in Figure 2, we measured the transcription start and end times as functions of the fraction of the nuclear cycle at which they occurred. As shown in Figure 3D, all transcription start times occurred at roughly 20% of the way through the nuclear cycle, while all transcription end times occurred at roughly 80% of the way through the nuclear cycle. Together, this implies that the time window during which transcription occurs comprises the middle 60% of the nuclear cycle across all measured temperatures, a result comparable to previous measurements performed at 25°C (Farrell and O’Farrell (2013)).

Having determined the consistent scaling of the transcriptional time window, we turned our attention to the RNA polymerase dwell time on the gene. Figure 3E shows the calculated dwell times that were extracted through the trapezoidal fitting described in the methodology behind Figure 3A. We calculated a dwell time for each position along the anterior-posterior axis in each embryo as part of the trapezoidal model fitting and averaged these values over multiple embryos. We found some variation in dwell time as a function of position along the embryo, though standard errors were large compared to parameter estimates (Figure S3). Following previous work indicating that *hunchback* dwell times were largely insensitive to position along the embryo between 20% and 40% of the embryo length (Liu et al. (2021)), we averaged the positionally-dependent dwell times over the same spatial window and over multiple embryos, to obtain the dwell time as a function of temperature alone (Figure 3E). Calculations of the dwell times normalized by the total cycle durations (Figure 3F) echoed our observations for the transcriptional start and end times: each RNA polymerase molecule takes approximately 20% of the nuclear cycle to elongate an mRNA molecule and terminate transcription irrespective of temperature.

We conclude that the timing of transcription—when it starts and ends during the nuclear cycle–as well as the dwell times of RNA polymerases on the gene— which capture both transcriptional elongation and termination—are temperature independent when measured in temporal units normalized by the temperature-specific nuclear cycle durations. To understand how these relationships to this normalized developmental time affect the amount of mRNA produced, and therefore the consequences of this temperature scaling for the regulatory program downstream of *hunchback*, we must also consider how the rate of transcriptional initiation depends on this environmental condition.

### Compensatory scaling of transcriptional time window and RNA polymerase loading rate lead to temperature-independent spatial patterns of produced mRNA

We next investigated how the mean rate of RNA polymerase loading, *R*, varies with temperature using the trapezoidal model introduced in Figure 3. As shown in Figure 4A, we found that loading rates increase with increasing temperature across the anterior half of the embryo.

**Figure 4.**
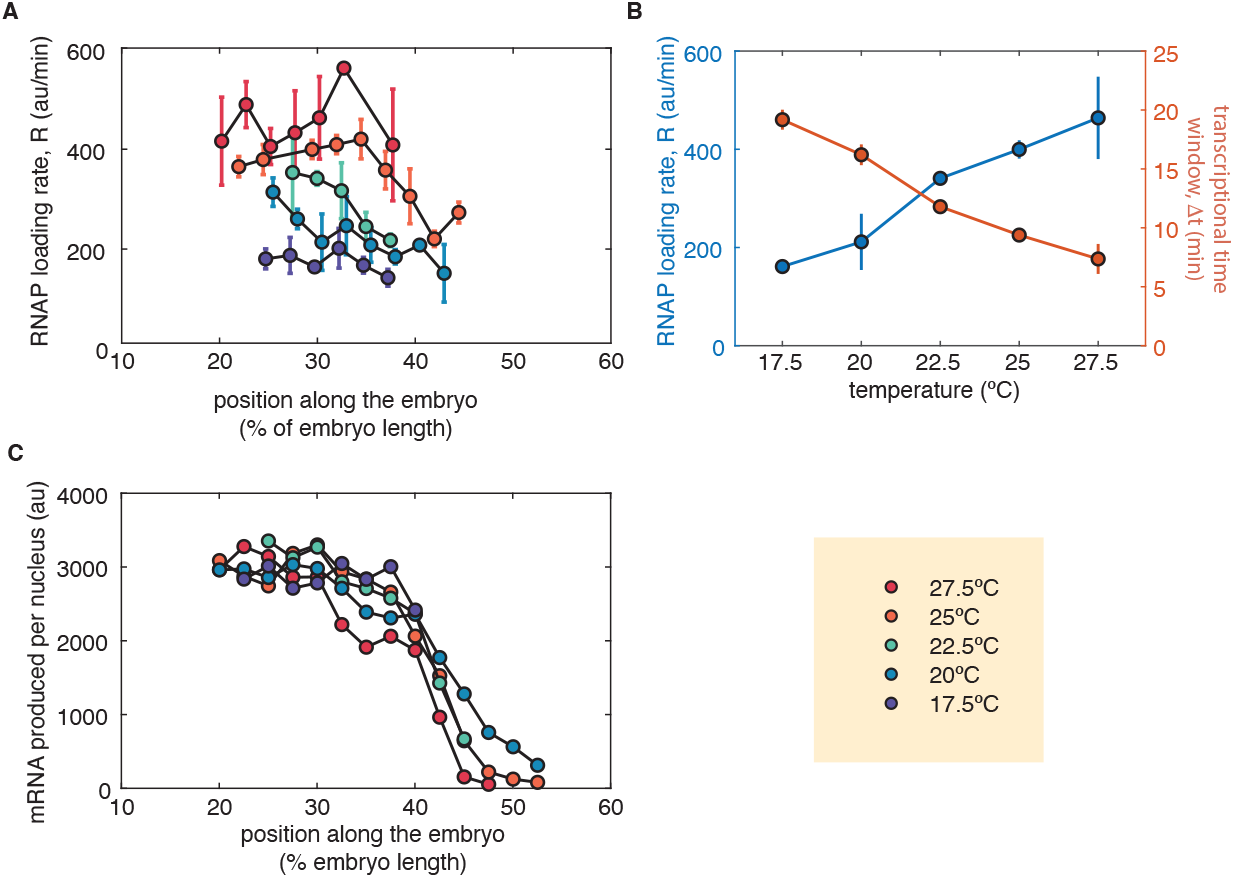
Compensatory scaling of initiation rate and transcriptional window duration lead to temperature-independent spatial patterns of produced mRNA. (A) Rate of RNA polymerase loading, *R*, as a function of the position along the embryo for different temperatures. Plotted values are the mean values calculated over parameters inferred from fits of the trapezoidal model to single embryo data, calculations included 2-5 embryos for each combination of temperature and anterior-posterior position, errorbars are standard errors over single embryo estimates. (B) RNA polymerase loading rate and duration of the transcriptional time window as a function of temperature measured at 30% along the anterior-posterior axis. (C) Total amount of mRNA produced along the embryo obtained by multiplying the loading rate and transcriptional time window. For errorbars (omitted here for legibility), see Figure S5

While Figures 3(D,F) and 4(A) revealed that the timing of transcription as well as the steps of the transcription cycle scale concomitantly with temperature, the quantity that ultimately matters for dictating regulatory programs downstream from *hunchback* is the pattern of mRNA production driven by the transcription cycle. At each temperature, this total amount of mRNA molecules produced can be readily calculated by multiplying the mean rate of transcription initiation by the duration of the transcriptional time window, as defined in Figure 3A.

Figure 4B shows the scaling of the mean transcription rate and the duration of the transcriptional time window at 30% of the embryo length as a function of temperature. The anti-correlation between these two magnitudes suggests that their product—resulting in the total amount of mRNA produced—might be independent of temperature. Indeed, when we calculate the total amount of mRNA produced as a function of position along the embryo for each temperature following the procedure described in Materials and Methods in order to estimate mRNA production at anterior-posterior positions where trapezoidal fitting failed due to low signal, we discovered, as shown in Figure 4C, that the the number of cycle 13 mRNA molecules is seemingly insensitive to temperature changes, such that the spatial profile of *hunchback* mRNA is unchanged. Similar results were obtained or positions with successful trapezoidal fits by, at each temperature and position along the embryo, multiplying these two magnitudes in order to obtain the total amount of mRNA produced (Figure S4). Thus, the transcription windows and polymerase loading rates scale in an inverse, compensatory fashion that ensures that the downstream number of produced *hunchback* mRNA molecules remains insensitive to temperature changes.

### Temperature scaling of the mean *hunchback* transcription rate is dictated by concomitant scaling of transcriptional bursting parameters

In the last few years it has been established that the expression of most developmental genes is characterized by transcriptional bursts, where the promoter stochastically transitions between an ON and an OFF state (Figure 5A, Lammers et al. (2020b)). This OFF state is seemingly distinct from the transcriptionally silent state observed before and after anaphase that ensues after *t*_*end*_ as defined in Figure 3A (Lammers et al. (2020a); Zhao et al. (2022)). Only when in the ON state does the promoter initiate transcription at a constant rate *r*, defined as the burst amplitude. The promoter switches between the OFF and ON states with a rate *k*_*on*_, typically defined as the burst frequency. Finally, the promoter transitions between the ON and OFF states with rate *k*_*off*_. We identify 1*/k*_*off*_ with the burst duration. Given these bursting parameters, the mean initiation rate *R* from the trapezoidal model described earlier, is then related to these bursting parameters by

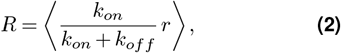

where the fraction accounts for the fraction of time the promoter spends in the ON state. Many transcription factors regulate the mean gene expression rate, *R*, by modulating the rates at which polymerases are loaded onto the gene in the ON state and the rates of promoter switching between the ON and OFF states (Lammers et al. (2020b)). Thus, transcriptional bursting provides an opportunity to determine whether, just like we saw for the mean transcription rate *R*, the temperature-scaling of the dynamics of the various bursting parameters that dictate the mean transcription rate are also coordinated and guided by the developmental clock.

**Figure 5.**
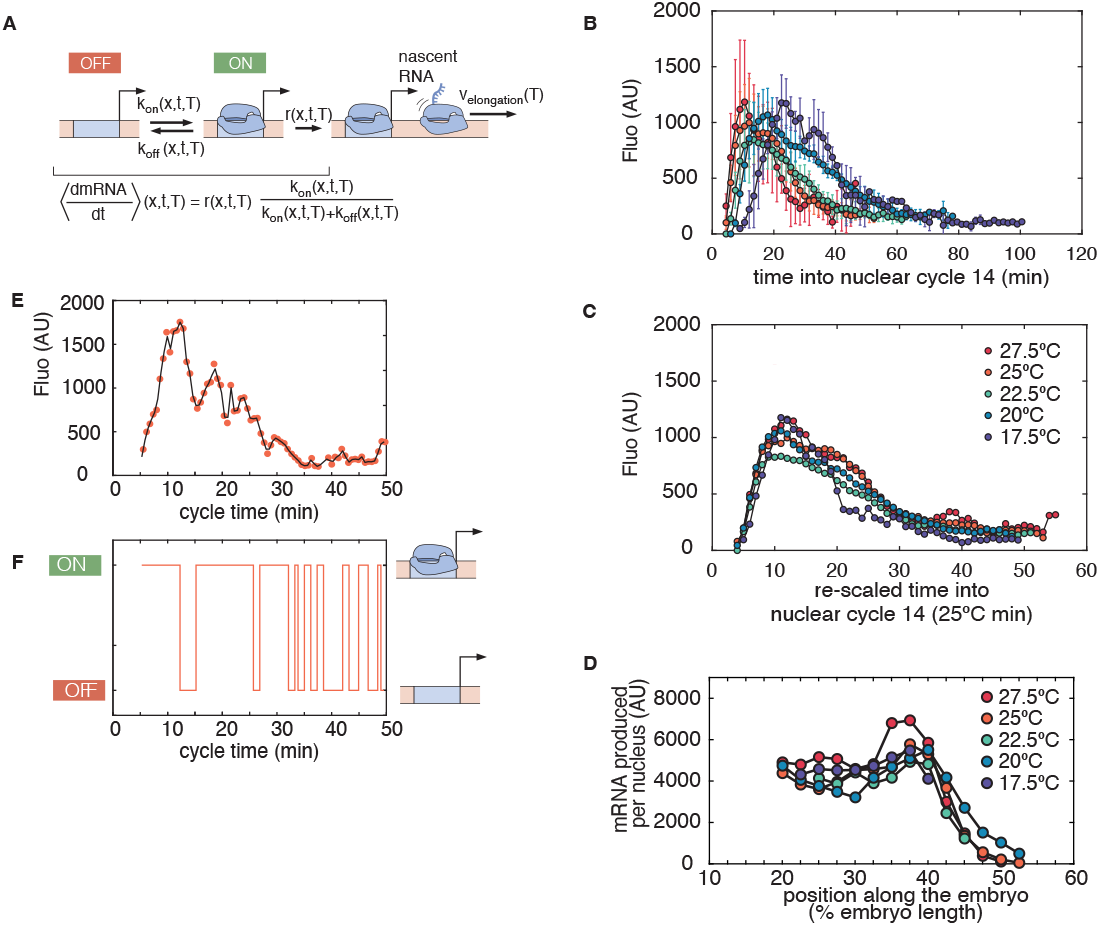
Transcriptional bursting in *hunchback* expression during nuclear cycle 14. (A) Two-state model of bursting of a single promoter defining burst frequency, duration and amplitude given by the parameters *k*_*on*_, 1*/k*_*off*_ and *r*, respectively. Note that these bursting parameters are functions of the position *x* (which determines the local concentration of input transcription factors), of time *t* and of the temperature *T*. (B) Mean MS2 fluorescence traces at 40% of the embryo length in nuclear cycle 14a. Note that the trapezoidal model introduced in Figure 3A no longer captures the transcriptional dynamics. (C) Time-rescaled version of the mean MS2 traces across temperatures at 40% of the embryo length in nuclear cycle 14.(D) Total mRNA produced in nuclear cycle 14 as a function of position along the embryo and temperature. (E) Representative single nucleus measurements at 40% embryo length for 25°C revealing that nuclei transcribe in “bursts” of activity. Black line is the inferred best fit trace from the cpHMM analysis. A windowed model was used for the inference, with 15 minute windows used at 25°C. (F) Inferred promoter state for the inferred best-fit trace showed in (E).

In *Drosophila melanogaster*, transcriptional bursts often last approximately 5 min and occur about every 5 min at room temperature, which is around 25°C (Lammers et al. (2020a)). As a result, each nucleus will experience only a handful—if any—of bursts during the short 17 min of nuclear cycle 13. The subsequent nuclear cycle, nuclear cycle 14, however, lasts much longer than nuclear cycle 13, stretching beyond 60 minutes at 25°C. This longer cycle typically presents multiple bursts, as shown in Figure 5E.

This nuclear cycle is also characterized by the degradation of the Bicoid gradient as well as significant increases in the expression of the other gap genes, which, together with *hunchback*, feed forward the positional information encoded by Bicoid and other gradients of maternally deposited proteins (Jaeger (2011)). In this context, *hunchback* exhibits a more dynamic pattern of transcriptional expression than it does in nuclear cycle 13, as reported by the mean MS2 fluorescence signals shown in Figure 5B. As seen in the figure, the number of actively transcribing RNA polymerase molecules sharply increases at the beginning of the nuclear cycle, followed by a slower decay. These observed dynamics clearly cannot be explained by the simple trapezoidal model introduced in Figure 3A, which is consistent with a constant mean rate of transcriptional initiation. Indeed, the data shown in Figure 5B imply that the rate of transcriptional initiation is modulated over time, and that the temporal dynamics of this modulation are temperature dependent.

Despite the fundamental difference in transcriptional initiation rate dynamics between nuclear cycles 13 and 14, an initial look at the cycle 14 data suggests cycle 14 transcriptional dynamics are governed by the same developmental clock that dictates cycle 13 dynamics. Because we were unable to capture the duration of cycle 14 across temperatures due to the loss of divisional synchronicity among nuclei and the movement of nuclei away from the embryonic surface, we used the cycle 13 durations across temperatures to map the times at distinct temperatures onto a common “developmental clock”. We used temporal measurements taken at 25°C as our reference (Table S1 and Figure S6). Specifically, we define the rescaled time into nuclear cycle 14, 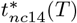 at a given temperature *T* as

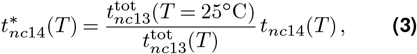

where 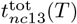 is the duration of nuclear cycle 13 at temperature *T* (referred to as *t*_*cycle*_ in previous sections) and *t*_*nc*14_(*T*) is the absolute time into nuclear cycle 14 at temperature *T*. Here, the rescaled time into nuclear cycle 14, 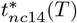, is the time at 25°C that we would expect to see the same expression patterns as we see at time *t*_*nc*14_(*T*) for temperature *T*, assuming that everything related to transcription is identical after accounting and correcting for differences in the total time over which expression occurs. Thus, 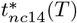 can be thought have as having units of “equivalent minutes at 25°C” or “25°C minutes”. This rescaled time is the cycle 14 analogue of the fractional progress through cycle 13 we used in our previous analysis of transcriptional dynamics in cycle 13. We also note that, taking equations 1 and 3 together implies that the scaling factor *α*(*T*) is given by

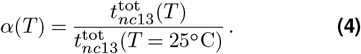

Using these mappings, we evaluate the MS2 signals across temperatures as functions of rescaled time. Just like in nuclear cycle 13 (Figure 2G), we discovered that the temperature-specific average MS2 fluorescence signals largely collapse onto a single curve within experimental error after re-scaling the time variables following Equation 3 and observing transcriptional activity as a function of “25°C minutes” (Figure 5C). While Figure 5C shows this collapse for measurements performed at 40% of the embryo length, this conclusion remains regardless of which position along the embryo is examined (Figure S7). Indeed, Figure 5D shows that this embryo-wide temperature collapse of the nuclear cycle 14 transcriptional dynamics leads to temperature-independent step-like pattern of produced mRNA across the embryo.

To gain insight into the collapse of average nuclear 14 transcriptional dynamics when plotted as a function of re-scaled time, we examined MS2 traces from single nuclei (examples are shown in Figure S8). Typical experimental traces, such as the one shown in Figure 5E, confirm that transcriptional activity occurs in stochastic “bursts” which appear to vary in amplitude, frequency and duration as the nuclear cycle progresses.

To measure the temporal and temperature dependence of the rates of promoter switching and polymerase loading in the ON state, we used a previously developed implementation of a compound-state Hidden Markov model (cpHMM, Lammers et al. (2020a)). The cpHMM allows us to uncover the underlying promoter state from the bursting MS2 signals extracted from individual nuclei, through which it infers values for *r, k*_*on*_, and *k*_*off*_. To allow for time-dependence in the rates of promoter-switching, we ran our inference using a sliding window of 15 minutes (see Methods and Materials for details.) The example experimental trace from Figure 5E is shown together with its “best fit”, which corresponds to the cpHMM-inferred promoter trajectory shown in Figure 5F.

To extract bursting parameter values, and their spatiotemporal and temperature dependence, we grouped traces by their position along the anterior-posterior axis and inferred bursting parameters as a function of time. Note that, for this analysis, we limited our inference to measurements taken between 22.5°C and 27.5°C due to computational constraints associated with longer dwell times at lower temperatures (see Methods and Materials for details).

Examining these parameters at 40% of the embryo length, we see that the polymerase loading rate, *r*, decreases over time across all observed temperatures (Figure 6A). These loading rates are also strongly temperature-dependent early in nuclear cycle 14, as was the case for the time-averaged loading rates we calculated for nuclear cycle 13. Similarly, burst durations decrease as a function of both time into cycle 14 and temperature (Figure 6B). However, across all temperatures, burst frequencies initially decrease before increasing again as the cycle progresses (Figure 6C). Though the overall temporal dynamics of burst frequency are similar for all temperatures assayed, the relationship between the burst frequencies and temperature is heterogeneous across time. Finally, the mean rate of transcription, *R*, obtained by combining our measurements of bursting parameters following Equation 2 is shown in Figure 6D. The figure reveals that the effect of the temporal and temperature modulation of each bursting parameter translates into a similar temporal and temperature dependence in the mean transcription rate.

**Figure 6.**
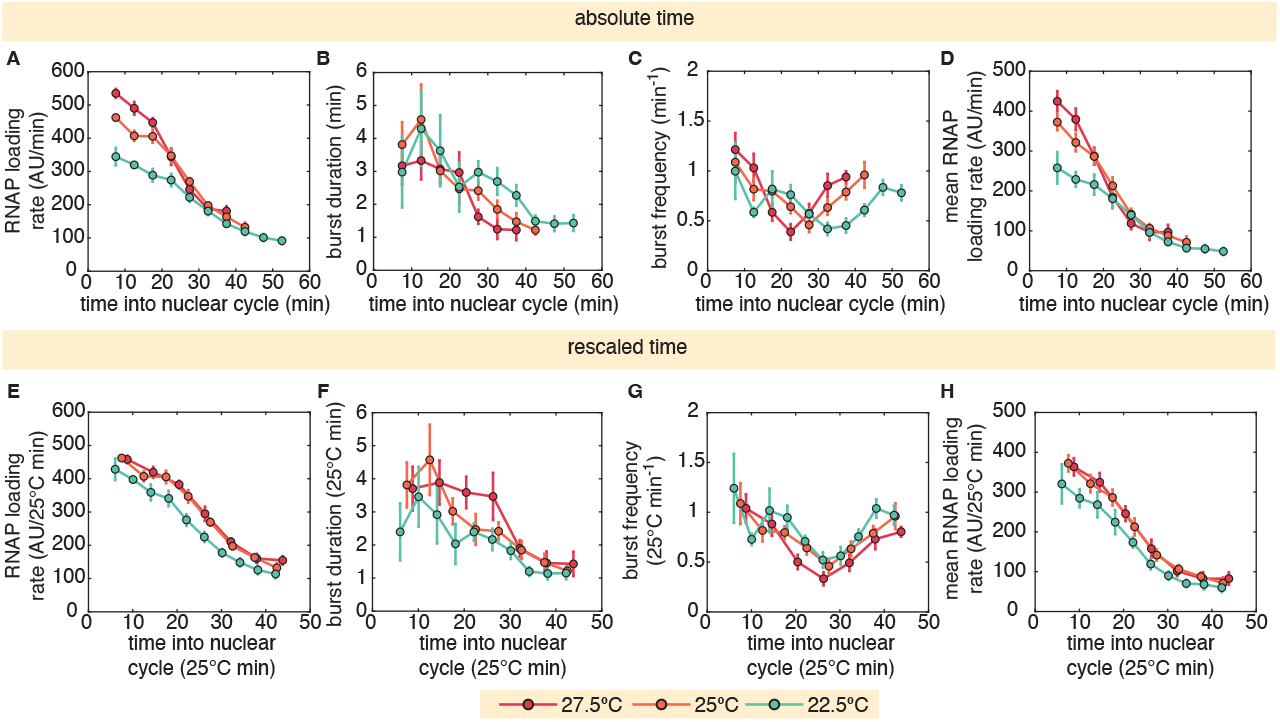
Temporal and temperature dependence of *hunchback* transcriptional bursting. (A-D) Temporal and temperature dependence inferred using cpHMM of (A) the RNAP loading rate *r*, (B) the burst duration 1*/k*_*off*_, and (C) the burst frequency *k*_*on*_. The temporal dependence of these parameters is similar in magnitude across temperatures, but the changes in these parameters with time are clearly temperature dependent. (D) Mean RNAP loading rate as a function of time calculated using Equation 2 recapitulates the characteristics of the instantaneous RNAP loading rates in (A). (E-H) Mapping the instantaneous RNAP loading rates to 25°C time (E) reveals that both the differences in the initial instantaneous loading rates and their variation with time are largely explained by the differences in the overall time of nuclear cycle 14. Similarly, differences in the time-dependence of both (F) burst durations and (G) frequencies are largely explained by mapping all three temperatures onto 25°C time. Finally, mapping the mean RNAP loading rates onto 25°C time (H) recapitulates the findings for the instantaneous loading rates in (E).

Like we have seen for all quantities dictating transcriptional dynamics throughout this work, we wondered whether discrepancies in the bursting parameter dynamics for different temperatures might be similarly explained by considering these quantities as functions of “25°C minutes” using equation 3. Using this method, we observed that temperature-specific discrepancies in both the scale and the timing of the polymerase loading rate, *r*, almost entirely disappear after we re-scale the time into “25°C minutes” (Figure 6E). We found similar results for the burst frequency and duration, shown in Figure 6F and G, respectively. It follows from our observations that much of the temperature-dependent distinction between the mean loading rate, *R*, is similarly explained by discrepancies in the overall temporal dynamics of the underlying bursting parameters, and that these dynamics are governed by the developmental clock.

## Discussion

Embryonic development is the result of a plethora of tightly coordinated biochemical reactions, the rates of which can potentially vary by different amounts as temperature fluctuates (Section). However, with one documented exception (Crapse et al. (2021)), the relative timing of the appearance of morphological landmarks (e.g., the time from fertilization to pole cell invagination in the fruit fly or the total time of embryogenesis) is independent of temperature (Kuntz and Eisen (2014)). These results suggest a model where a “developmental clock” sets a unique temperature scaling which maintains coordination of all processes underlying embryogenesis (Kuntz and Eisen (2014)), as described mathematically by the scaling relation in Equation 1.

Whether this unique temperature scaling operates beyond macroscopic morphological processes and ensures the coordination of the molecular processes underlying cellular decision making, however, remains an open question. Specifically, if the developmental clock model applies at the microscopic level, the relative rates of the processes of transcription and the cell cycle that are so fundamental to cellular fate specification should be temperature independent. Another possibility is that the rates of different microscopic processes happening in parallel change by different factors with temperature. Such uncoordinated changes in microscopic rates could lead, for example, to a temperature-dependent modulation of the spatiotemporal dynamics of mRNA patterns, and this modulation would then propagate through the gene regulatory network. In this case, feedback mechanisms could exist that correct for any potential deviations in the timing of the establishment of cell fate. Such compensatory mechanisms have been shown to exist at the transcriptional level to account for changes in the dosage of morphogen gradients (Liu and Ma (2011)).

Clearly, to shed light on the molecular origins of this developmental clock, it is necessary to go beyond measurements of the temperature scaling of the timing of appearance of coarse-grained morphological features to uncover the scaling of the biochemical processes that underlie these macroscopic changes. A recent work focusing on the scaling of the phases of the cell cycle in the early fly embryo found that, with the exception of prophase, all remaining stages of the cycle—interphase, prometaphase, metaphase, anaphase, telophase—displayed a comparable scaling with temperature, once again consistent with Equation 1 (Falahati et al. (2021)). However, until now, these sorts of studies had not been extended to the temperature scaling of any other molecular processes underlying embryonic development.

In this paper, we quantified the temperature scaling of the main processes of the transcription cycle—initiation, elongation and termination—as well as the timing of the start and end of transcription with respect to the cell cycle in living fruit fly embryos. Within experimental error, we found that all rates—or effective rates— follow the same quantitative temperature scaling *α*(*T*) as defined in Equation 1. Further, we zoomed into the mean rate of transcription and dissected it in terms of its underlying transcriptional bursting and the paremeters that dictate the dynamics of these bursts. Here, we found a rich spatiotemporal dependence of all bursting parameters—frequency, duration and amplitude. However, once again, the temperature scaling of these bursting parameters was quantitatively indistinguishable from each other, and from any of the other parameters we dissected in this work. Thus, we conclude that the developmental clock proposed to operate at the macroscopic level might also be at play in dictating the pace of the molecular processes underlying gene regulatory pattern dynamics and the subsequent cellular decision making.

The molecular nature of the developmental clock ensuring the coordination of both microscopic and macroscopic developmental processes remains unknown. One possibility is that evolution has selected for all underlying molecular reactions to scale in the same fashion with temperature. This explanation, while possible, seems at odds with the range of activation energies for enzymatic reactions measured throughout biology (Section and Lepock (2005)). Another possibility is that there is an actual molecular player that constitutes a rate-limiting step for the plethora of reactions that lead to cellular decision making, and that therefore could set the pace of these reactions. For example, ATP consumption could simultaneously dictate the rate of multiple reactions, as has been seen in the cyanobacterial cyrcadian clock (Terauchi et al. (2007)). Previous measurements of the temperature sensitivity of the the mitotic cycle explored the existence of a kinase-based developmental clock (Falahati et al. (2021)). However, the entry into prophase was prolonged relative to other cell cycle transitions at lower developmental temperatures, suggesting that multiple kinases—and not just a unique molecular player—must be involved in pacing development when considered at this level of temporal granularity.

Perhaps one of the most interesting aspects of the universal scaling relationships we extended with respect to previous works (Kuntz and Eisen (2014); Crapse et al. (2021); Falahati et al. (2021)) is that, due to the compensated increase in transcriptional initiation and decrease in length of the nuclear cycle with increasing temperature, the total amount of mRNA produced in one cycle is temperature-independent. Of course, it is the cytoplasmic mRNA concentration, and not the total amount of produced mRNA, that dictates the downstream protein pattern that feeds back into the gene regulatory network to ensure proper developmental outcomes. To predict the spatiotemporal dynamics of the cytoplasmic concentration of mRNA and protein, it is necessary to determine the temperature scaling of the rate of protein translation as well as the rates of mRNA and protein degradation and their diffusion constants. The combined temperature scaling of all these molecular processes, if not matched as in the processes dissected throughout this work, could have significant effects on the spatiotemporal dynamics of the cytoplasmic protein patterns that drive the developmental program.

Our current experimental design does not afford access to the molecular parameters necessary to predict protein concentration dynamics from the MS2 signal reporting on the instantaneous number of RNAP molecules actively transcribing the gene. However, various approaches do exist for measuring, with high spatiotemporal resolution, the protein diffusion rate (Abu-Arish et al. (2010); Athilingam et al. (2023)), the protein degradation rate (Bothma et al. (2018)), and the protein translation rate (Vinter et al. (2021); Dufourt et al. (2021)) in fly embryos. To our knowledge, measurements of the mRNA degradation rate are not yet possible in single cells of living embryos, but measurements in precisely timed embryos combined with RNA sequencing (Thomsen et al. (2010); Little et al. (2013); Forbes Beadle et al. (2023)) can shed light on the embryo-averaged temporal dependence of this magnitude. Further, while mRNA diffusion rates have not been measured in flies, they have been quantified in single cultured cells (Yan et al. (2016)). In principle, this same technology could be deployed in the context of the fruit fly to measure temperature-dependent mRNA diffusion rates.

It is important to note that, so far, the concept of a developmental clock that sets the pace of development regardless of temperature has been a useful model to understand the temperature-scaling phenomenology observed (Kuntz and Eisen (2014); Crapse et al. (2021); Falahati et al. (2021)). Indeed, this model seems to hold at both the macroscopic and microscopic levels. Yet, a few exceptions to this the unified scaling of the molecular clock have already been uncovered (Crapse et al. (2021); Falahati et al. (2021)). We speculate that it is these exceptions that have the potential to uncover whether the concept of the developmental clock has a molecular counterpart. Thus, we propose that parsing the temperature dependence of an increasing swath of molecular processes and revealing whether their temperature scaling remains coordinated with respect to each other or not—as was recently uncovered in the context of the cell cycle (Falahati et al. (2021))—-might shed clues on the molecular nature—and limits—of the developmental clock.

Finally, and regardless of the molecular origins of the developmental clock, we are excited to use temperature as a knob for the *in vivo* biochemical dissection of the molecular processes that drive cellular decision making. Specifically, quantifying the dynamics of the processes of the central dogma as temperature is varied could reveal the existence of intermediate molecular steps and their kinetics, as was done, for example, in the *in vitro* context to dissect bacterial transcription (Record et al. (1996)).

## Methods

### Reporter construct

This work used fly lines developed in ref. (Bothma et al. (2015)). The *hunchback* reporter construct contains 24 repeats of the MS2 stem loop sequence (often referred to as the MS2 cassette) inserted in the 5’ UTR of the *hunchback* reporter gene. The endogenous *hunchback* coding sequence was replaced with that of the yellow-kanamycin reporter gene. The reporter gene was integrated at PBac{y[+]-attP-3B}VK00002 on the second chromosome by injecting strain BDSC 9723.

### Temperature control

The temperature during imaging was primarily controlled using a stage-top water-jacket incubator made by Okolab (H101-LG, Napoli, Italy) with water circulating from a static temperature water bath (LAUDA ECO RE 415 S 4L Water Bath circulator with ECO Silver Control Head, LAUDA-Brinkmann, Marlton, NJ) to control the temperature inside of the chamber. The water-jacket incubator is comprised of a main body that screws into the microscope stage and holds the sample, and a lid, both of which are heated and cooled by the flow of water from the thermostatic bath through interior. The sample temperature is therefore set by the heat and cool emanating from the body of the chamber. An objective collar with similar channels for water flow is used to directly control the temperature of the objective, which is in direct thermal contact with the slide-mounted sample within the chamber. Additionally, we controlled the temperature of the room that housed the microscope as a secondary measure for minimizing any potential gradients across the sample holder. Insulating curtains were used to thermally isolate this room to the fullest extent possible.

A fine gage thermocouple inserted through holes in the back of the membranes on which embryos were mounted was used to measure temperatures between the oxygen permeable membrane and the coverslips. Because the thermocouple’s diameter was larger than that of the fly embryos (*∼*130*μm*), temperature could not be monitored during the imaging experiments without disrupting the mounted embryos. We therefore calibrated the relationship between the room temperature, bath temperature and sample temperature before beginning embryo imaging experiments, confirming that slide temperatures were stable over periods of many hours and that any temperature gradients across slides were minimized. Additionally, at the end of each imaging session when data was no longer being collected for the embryos, we confirmed that temperatures between the coverslips and membranes were within 0.2°C of the intended sample temperature.

### Embryo preparation and data collection

Sample preparation followed procedures described in previous work (Garcia et al. (2013); Bothma et al. (2015)). Female virgins of yw;HisRFP;MCP-GFP (MCP, MS2 Coat Protein) genotype were crossed with males bearing the *hunchback* reporter gene. Collection chambers were kept in incubators at 25°C for several days before embryo collection began. Collection plates were changed every thirty minutes for at least an hour before beginning collections for experiments to ensure that all embryos were newly fertilized.

After two or more rounds of clearing, embryos were collected in thirty minute intervals. Collected embryos were dechorionated using bleach and mounted in halocarbon oil 27 between a semipermeable membrane (Lumox film; Starstedt) and a coverslip, with care taken to limit the time embryos spent outside of the temperature chamber after the initial collection window to no more than fifteen minutes. Embryos were then incubated within the sample chamber for 2-6 hours depending on the experiment temperature until cellularization was well under way in nuclear cycle 14 and the cellular migration was beginning to occur.

Data was collected using a Leica SP8 laser scanning confocal microscope with a 63x/1.40 oil objective. Image resolution was 856 × 856 pixels, with a pixel size of 108 nm. At each time point, a stack of 21 images separated by 500 nm was collected. Image stacks were collected at a time resolution of *∼*37*s*. The MCP-GFP and His-RFP were excited with wavelengths of 488 nm and 556 nm, respectively, using a White Light Laser. The average laser power (measured using a 10x air objective) used for imaging the embryos was 35*μ*W at 488 nm and 25*μ*W at 556 nm. Two Hybrid Detectors (HyD) were used to detect the fluorescence signals over windows of 498-556 nm and 566-724 nm.

### Image Analysis

Image analysis of live embryo movies was performed based on the protocol in (Garcia et al. (2013)) with modifications to this protocol, including the use of the Trainable Weka Segmentation plugin for FIJI using the FastRandomForest algorithm as described in (Lammers et al. (2020a)).

### Measurement of nuclear cycle length

As it was previously shown that cell cycle duration is strongly determined by developmental temperature (Kuntz and Eisen (2014); Falahati et al. (2021)), we used nuclear cycle duration to confirm that embryogenesis was proceeding at a rate in reasonable agreement with our expectations for the developmental temperature. To extract the mean nuclear cycle duration for each embryo movie, we maximum projected the Histone-RFP slices for each time point. Nuclei were segmented and tracked over multiple nuclear cycles. For each nucleus, the start of the nuclear cycle was determined by the first frame in which the nucleus appeared to be in anaphase and was distinguishable from its sister cell. For each embryo, nuclear cycle durations were calculated by averaging the length of the division cycle over all individual nuclei that were observed in every frame between successive anaphases. Between four and thirteen embryos, each with between eighty and one hundred persistent cycle 13 nuclei in the field of view, were observed at each temperature. Single-embryo estimates of nuclear cycle duration were than averaged to calculate temperature-specific nuclear cycle durations for each developmental temperature.

### MS2 signal quantification

Nuclear traces were aligned so that, for each fluorescence time trace, *t* = 0 was the first frame in which the associated nucleus was observed in anaphase. Nuclear traces were then resampled to 60 second resolution and averaged together over a spatial window corresponding to 2.5% of the anterior-posterior embryo length. For each temperature, AP-dependent fluorescence measurements were then averaged over embryos to construct a temperature-specific mean expression level as a function of time into the nuclear cycle and position along the anterior-posterior axis.

### Correcting for temperature-dependence in GFP fluorescence intensity

The fluorescence intensity of GFP is temperature dependent. We used previous measurements of the temperature dependence of GFP (Zhang et al. (2009)) to re-scale our fluorescence intensity measurements so that we could make direct comparisons between MS2 signals captured at different temperatures. We extrapolated fluorescence coefficients based on the observed linear relationship of fluorescence intensity and temperature over the range between 17.5°C and 27.5°C. We the calculated fluorescence scaling coefficients for each of our observed temperatures, once again using 25°C as the reference temperature and therefore setting its scaling factor to 1. The remaining fluorescence scaling coefficients are summarized in Table S2 and plotted in Figure S9. We then multiplied the fluorescence signals from single nuclei by the fluorescence intensity scaling coefficient for the relevant temperature, allowing for comparison between data from different temperatures.

### Trapezoid parameter extraction

Trapezoidal fluorescence intensities were fit using a custom fitting function. Mean traces are smoothed and the smoothed traces are used to determine the times into the nuclear cycle corresponding to the beginning and end of the plateau portion of the trapezoidal signal. The positive and negative slopes of the plateau are fit independently to determine the start and end times of the transcriptional window, and the loading and unloading rates of polymerases. The remaining time points within the plateau are then fit to determine the plateau height, from which the dwell time through the intersections of the different piecewise components of the trapezoidal function.

### Calculating total mRNA production

To calculate the total mRNA produced in fluorescence units, we related the fluorescence time trace to the amount of mRNA produced. The mean fluorescence trace is the sum of the individual fluorescence signals produced by single polymerases transcribing the reporter. Each polymerase produces a square-wave fluorescence signal with a time equal to the polymerase’s elongation time and an amplitude equal to the fluorescence of a single saturated 24xMS2 stem loop cassette. Equivalently, the integrated fluorescence of a single polymerase square-wave signal over time divided by the elongation time is equivalent the instantaneous fluorescence of a single polymerase transcribing the reporter. The cumulative signal of these individual polymerase fluorescence pulses added together gives the trapezoidal fluorescence signal observed from the reporter (Figure 3B). Since every polymerase transcribing at the same temperature has approximately the same elongation time, the mean number of polymerases transcribing is proportional to the integral of mean fluorescence divided by the elongation time.

The number of mRNA molecules produced per nucleus were calculated by dividing the mean integrated fluorescence at a given AP and temperature by the elongation time at that temperature. Error bars were calculated by propagating the standard error over embryos of the integrated fluorescence and the elongation time.

### Hidden Markov Model and Elongation Times

See Lammers et al. (2020a).

---

**Figure S1.**
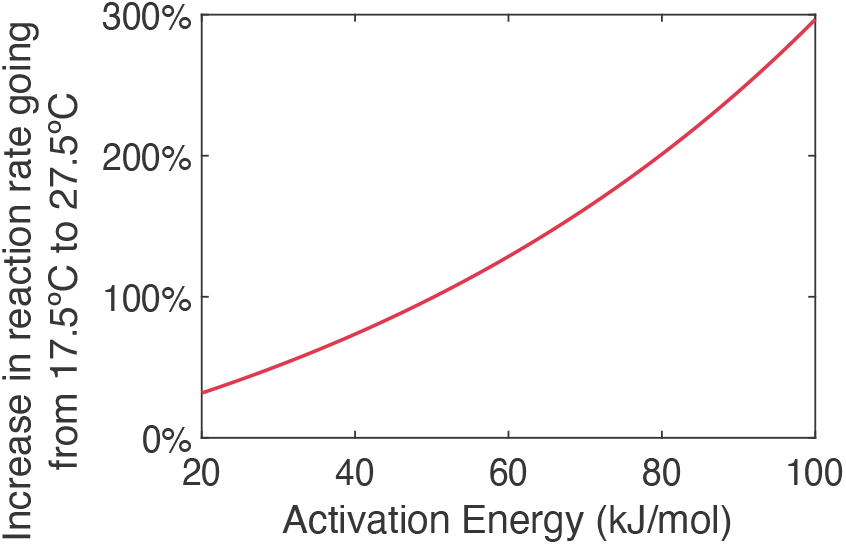
Temperature dependence of molecular rates as a function of activation energy. Arrhenius-predicted percentage increase in rates of the same reaction going from 17.5 °C to 27.5°C for activation energies ranging between 20 and 100 kJ/mol.

**Table S1.** Summary of time-scaling coefficients for all developmental temperatures.

| Development temperature (°C) | Cycle 13 duration (min) | Time-scaling coefficient |
| --- | --- | --- |
| 27.5 | 14.57 | 0.86 |
| 25 | 17.03 | 1.0 |
| 22.5 | 21.17 | 1.24 |
| 20 | 27.34 | 1.61 |
| 17.5 | 34.86 | 2.05 |

## Supplementary Information

### Arrhenius calculation

In the simplest model, which has been repeatedly confirmed experimentally, first-order biochemical reactions, characterized by a rate *k*, scale through the so-called Arrhenius relation given by

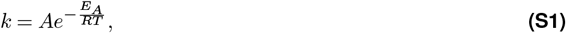

where *A* is a temperature-independent prefactor, *E*_*A*_ is the activation energy of the reaction, *R* is the gas constant, and *T* is the temperature in Kelvin. For most enzymatic reactions, activation energies fall in a range between 20 and 100 kJ/mol (Lepock (2005)). Consider, for example, the the 10°C range—from 17.5°C to 27.5°C—over which *Drosophila melanogaster* lives and develops healthily (Kuntz and Eisen (2014)), and that we will entertain throughout this work. The rate of a process with an underlying activation energy of 20 kJ/mol will increase by 40% when changing the temperature between 17.5°C and 27.5°C, while the rate of a process with an activation energy of 100 kJ/mol will increase by 300% over the same temperature range (Figure S1). Thus, the rates of the biochemical processes dictating embryonic development might be expected to vary by drastically different amounts as temperature changes, presenting challenges to ensuring a robust, coordinated and reproducible outcome for the precisely orchestrated process of animal development.

**Table S2.** Summary of fluorescence scaling coefficients for all developmental temperatures.

| Development temperature (°C) | Fluorescence scaling coefficient |
| --- | --- |
| 27.5 | 1.087 |
| 25 | 1.000 |
| 22.5 | 0.9259 |
| 20 | 0.8620 |
| 17.5 | 0.8063 |

**Figure S2.**
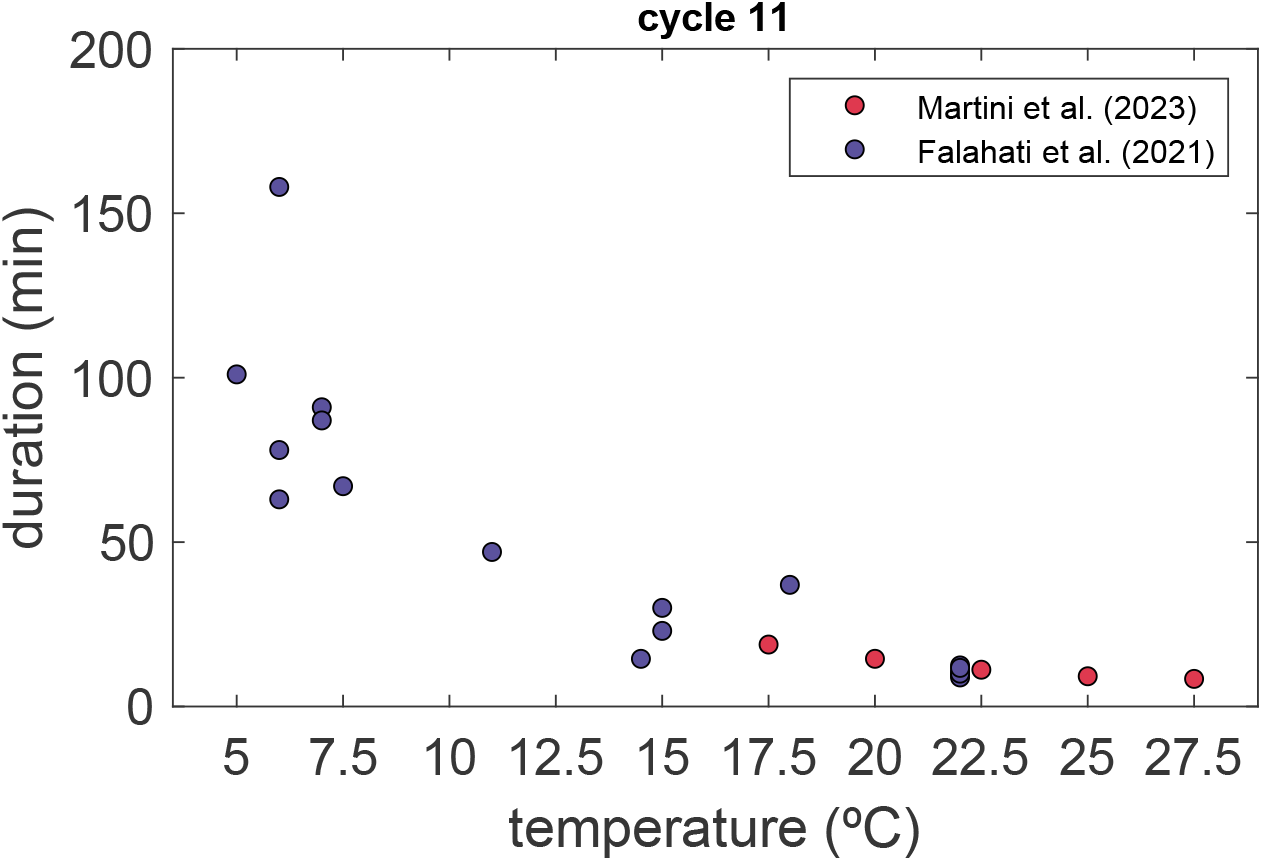
Comparison of measured nuclear cycle 11 durations and nuclear cycle 11 durations reported by Falahati et al. (2021).

**Figure S3.**
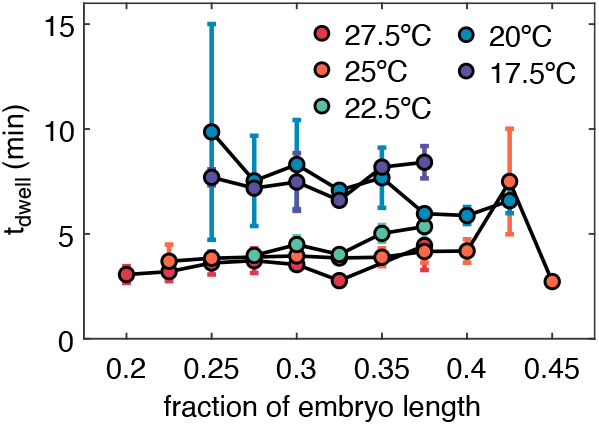
RNA polymerase dwell times along the embryo. Dwell time *t*_*dwell*_ as a function of position along the anterior-posterior axis. Dwell times were inferred using the trapezoidal fitting introduced in Figure 3(A).

**Figure S4.**
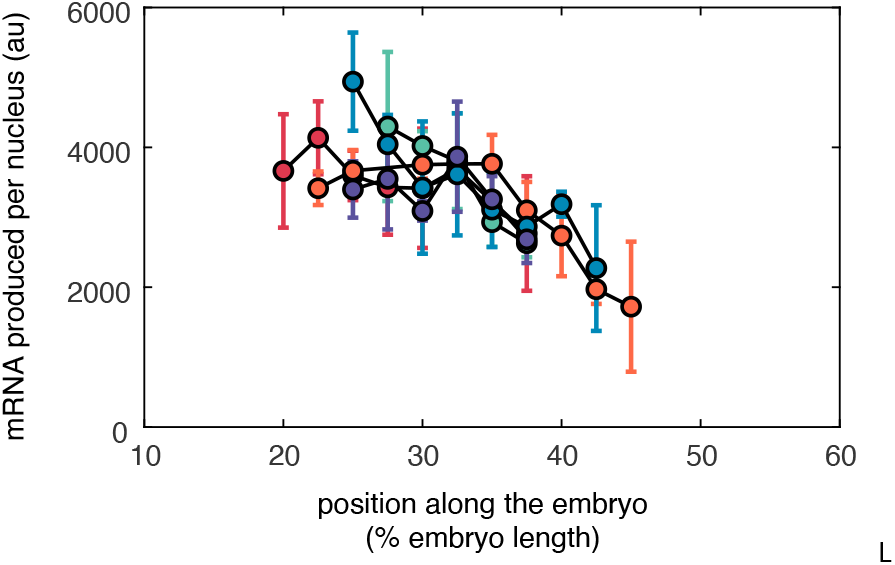
Total mRNA produced in nuclear cycle 13, calculated by multiplying the mean initiation rate by the transcription window.

**Figure S5.**
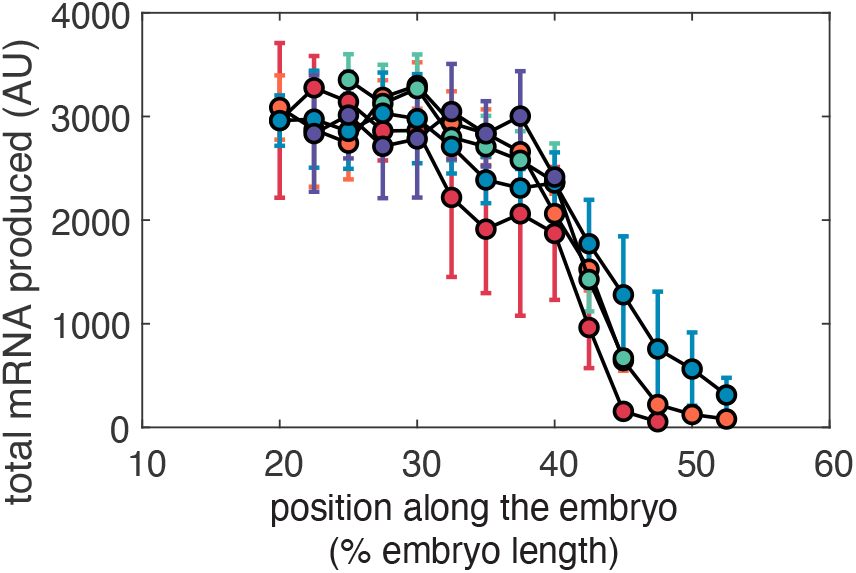
Total mRNA produced in cycle 13 as a function of temperature and position along the anterior-posterior axix of the embryo (as shown without errorbars in Figure 3E).

**Figure S6.**
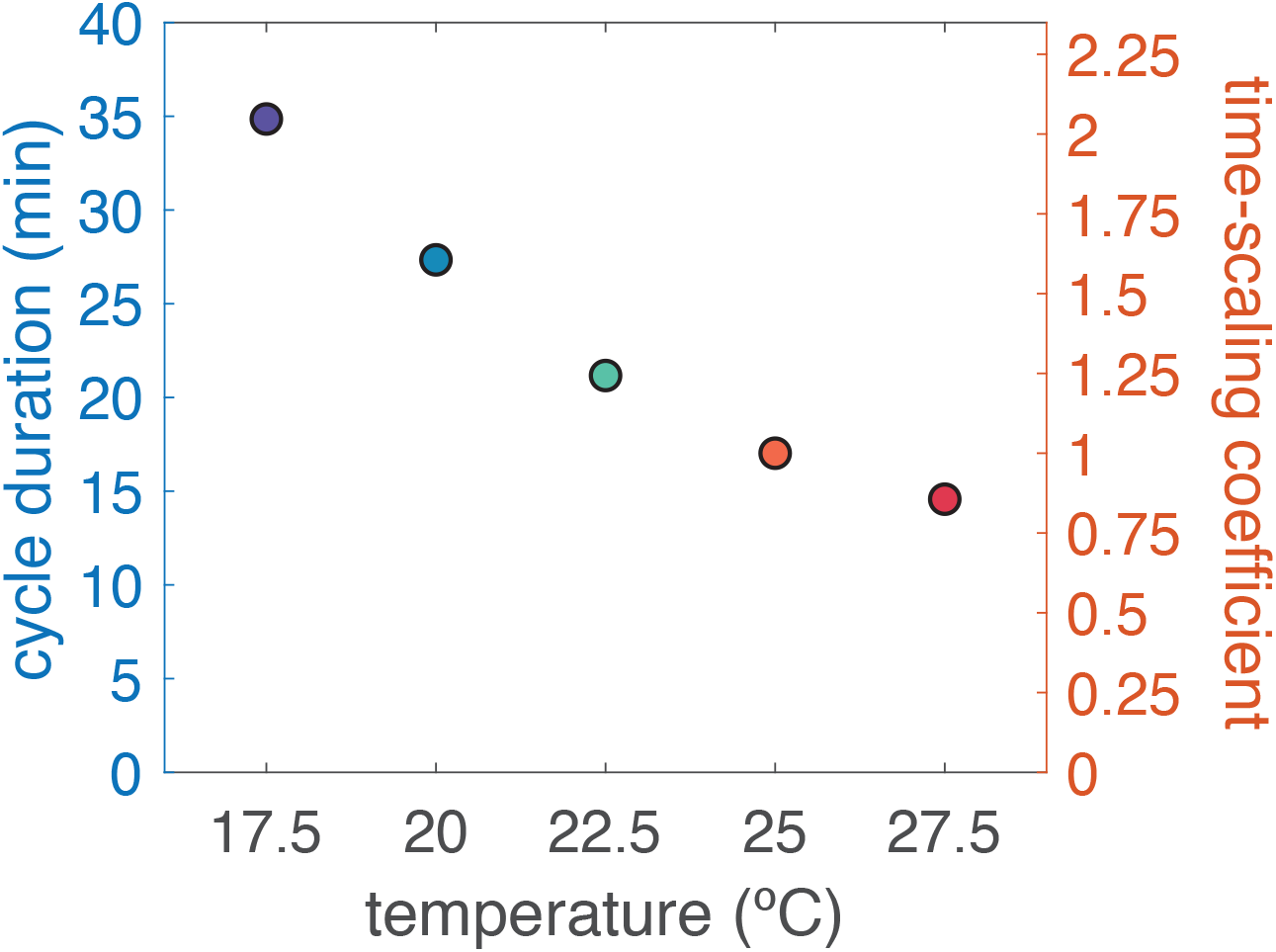
Nuclear cycle 13 durations with time in minutes shown on the left axis and normalized time-scaling coefficients shown on the right axis.

**Figure S7.**
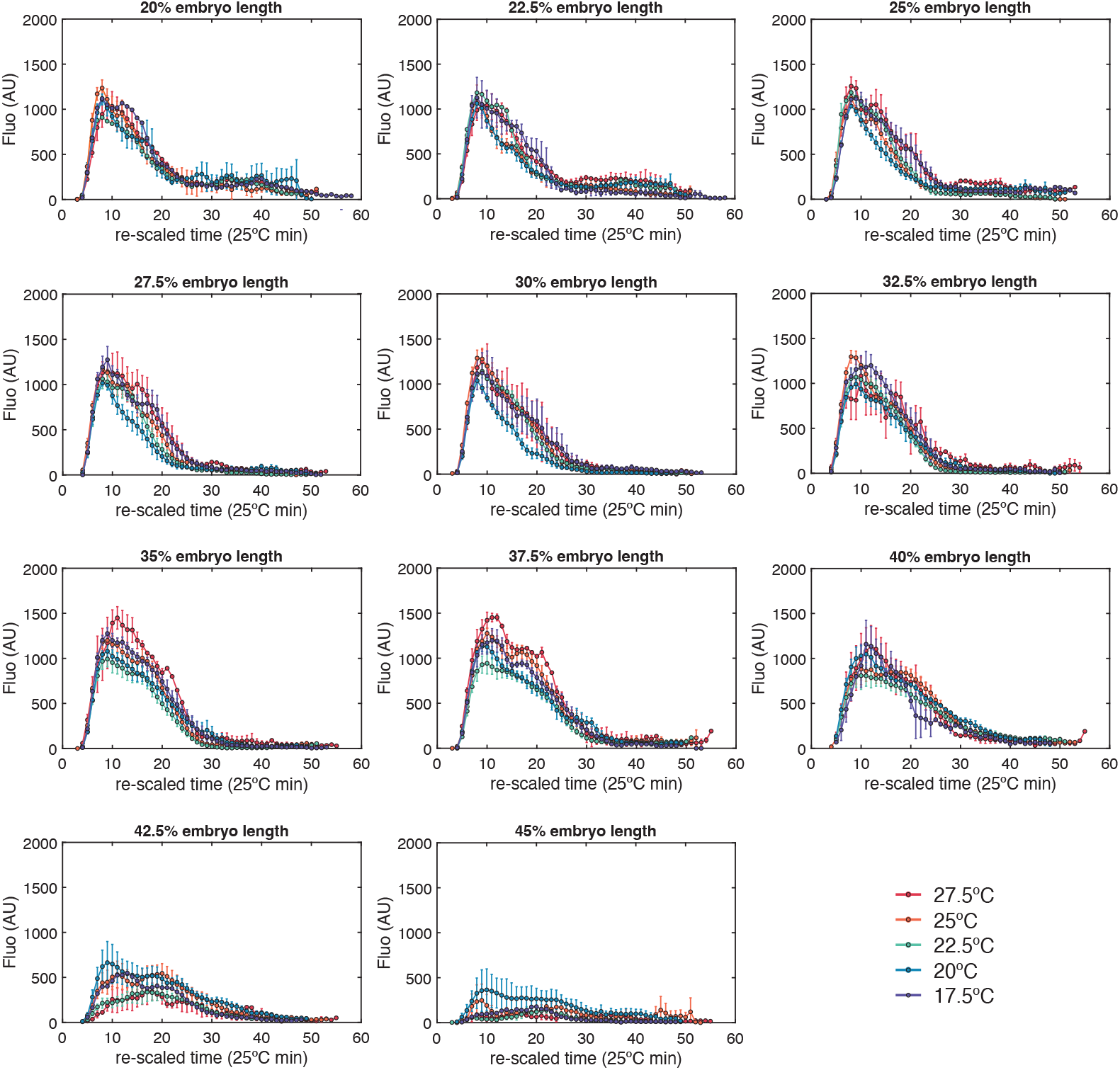
Nuclear cycle 14 mean normalized profiles.

**Figure S8.**
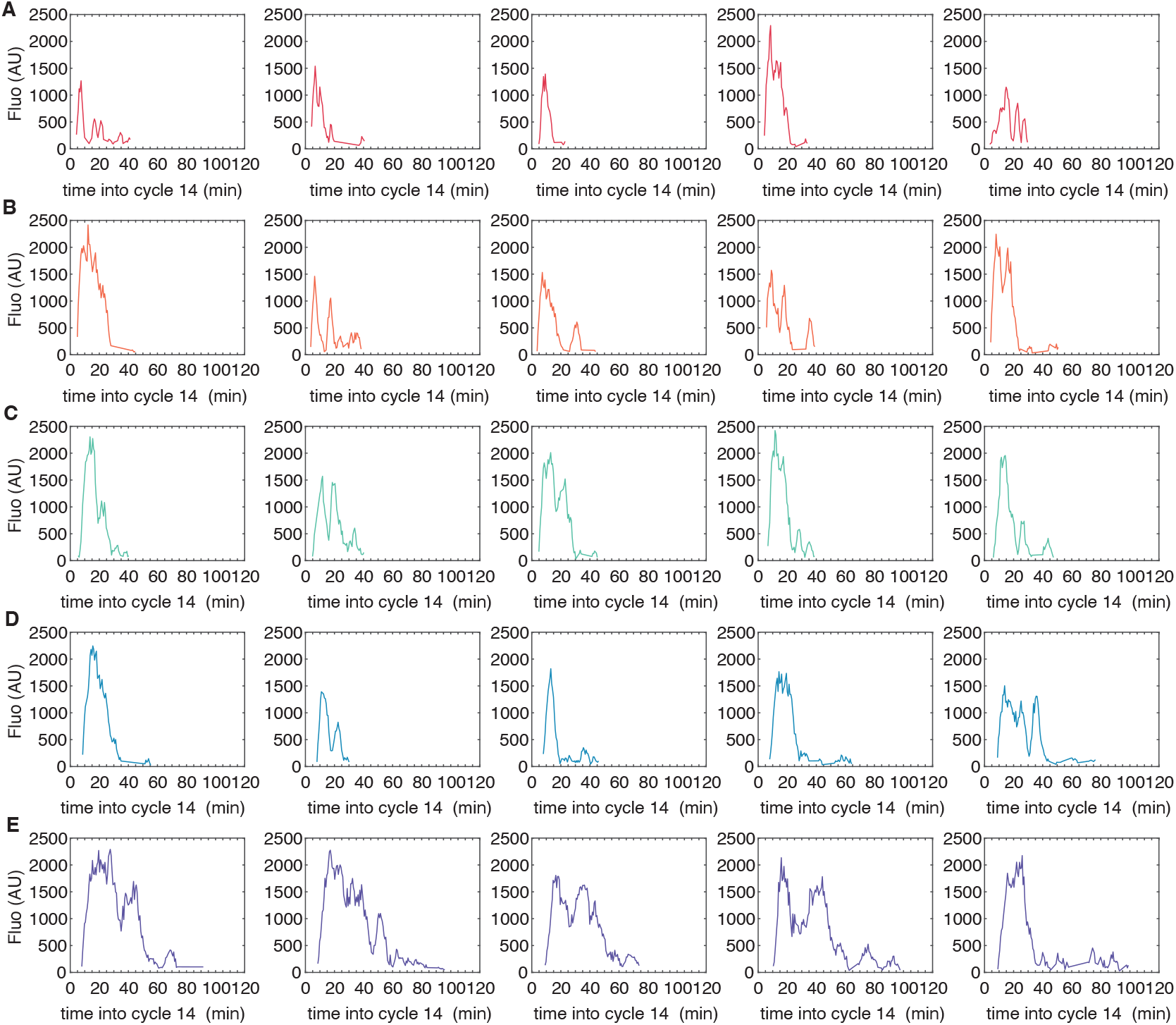
Representative traces for MS2 reporter signals from single nuclei in cycle 14 at 30% AP position for embryos developing at 27.5°C (A), 25°C (B), 22.5°C (C), 20°C (D), and 17.5°C (E).

**Figure S9.**
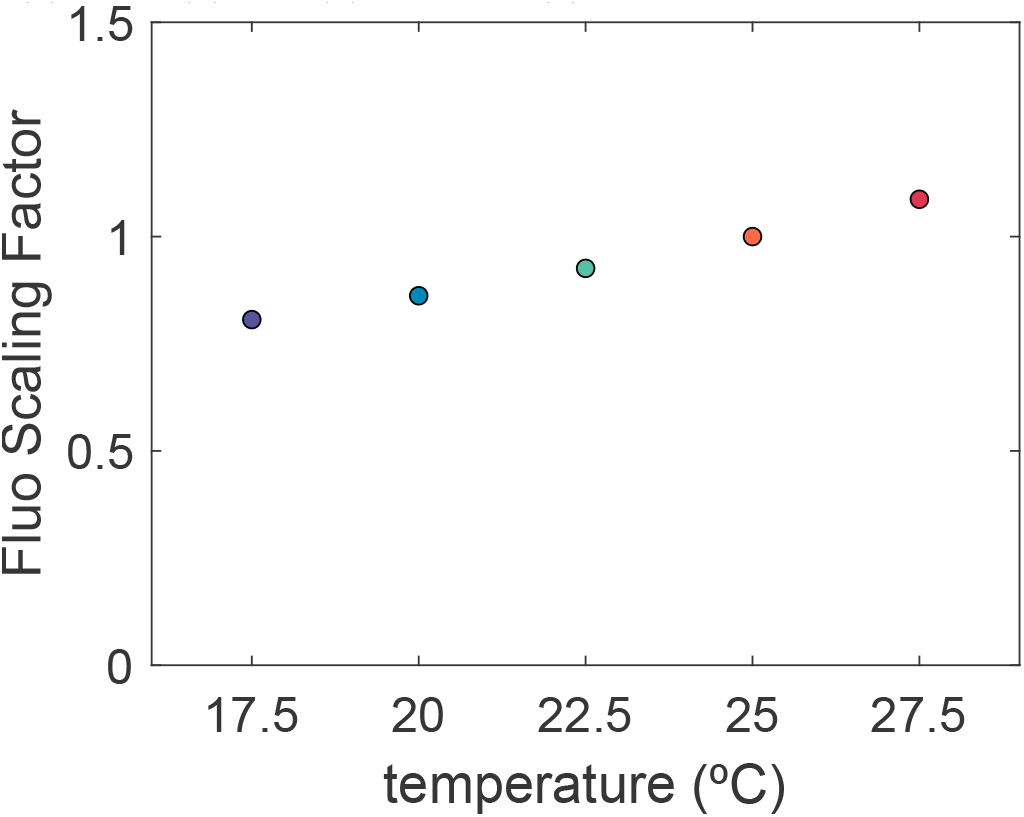
Fluorescence scaling factors used to correct for temperature-dependent differences in GFP fluorescence intensities.

